# Coffee polyphenols prevent cognitive dysfunction and suppress amyloid β plaques in APP/PS2 transgenic mouse

**DOI:** 10.1101/347963

**Authors:** Keiko Ishida, Masaki Yamamoto, Koichi Misawa, Noriyasu Ota, Akira Shimotoyodome

**Affiliations:** Biological Science Laboratories, Kao Corporation, Haga-gun, Tochigi, Japan; Health Care Food Research Laboratories, Kao Corporation, Sumida, Tokyo, Japan

**Author notes:** Corresponding author: Keiko Ishida, Ph.D. E-mail address (KI).

## Abstract

Epidemiological studies have found that habitual coffee consumption may reduce the risk of Alzheimer’s disease. Coffee contains numerous phenolic compounds (coffee polyphenols) such as chlorogenic acids. However, evidence demonstrating the contribution of chlorogenic acids in preventing cognitive dysfunction induced by Alzheimer’s disease is limited. In this study, we investigated the effect of chlorogenic acids on prevention of cognitive dysfunction in APP/PS2 transgenic mouse model of Alzheimer’s disease. Five-week-old APP/PS2 mice were administered a diet supplemented with coffee polyphenols daily for 5 months. The memory and cognitive function of mice was determined using the novel object recognition test, the Morris water maze test, and the step-through passive avoidance test. We found that chronic treatment with coffee polyphenols prevented cognitive dysfunction and significantly reduced hippocampal Aβ deposition. We then determined the effect of 5-caffeoylquinic acid, one of the primary components of coffee polyphenols, on Aβ formation. 5-Caffeoylquinic acid did not inhibit Aβ fibrillation, but degraded Aβ fibrils in a dose-dependent manner. In conclusion, these results demonstrate that coffee polyphenols prevented cognitive deficits and alleviated Aβ plaque deposition via disaggregation of Aβ in APP/PS2 mouse.

## Introduction

Alzheimer’s disease (AD), a neurodegenerative disease characterized by memory and cognitive dysfunction, is a worldwide public health problem. Its pathological features include the accumulation of amyloid β (Aβ) peptides (amyloid plaque) and aggregation of neurofibrillary tangles, which play causal and central roles in AD progression [1, 2]. Aβ accumulation is due to an imbalance between its synthesis and clearance. Aβ is generated through proteolytic processing that involves two types of proteases, namely, β- and γ-secretase [3]. A transmembrane aspartic protease called β-site APP cleaving enzyme (BACE1) is responsible for β-secretase activity. Although BACE1 cleaves APP and releases the soluble domain out of the cell, Aβ body remains attached to the C-terminal fragment (C99) bound to the cell membrane [4]. Furthermore, C99 undergoes cleavage by γ-secretase and releases Aβ isoforms such as Aβ 1–40 and Aβ 1–42 [5]. Conversely in the nonamyloidogenic pathway, α-secretase cleaves amyloid-β precursor protein (APP) within the Aβ domain and produces soluble APP-α, thus preventing Aβ. To date, no curative treatment for AD has been reported; most therapeutic approaches transiently relieve symptoms and do not improve or inhibit disease progression. The dominantly inherited Alzheimer’s network study, which observes changes in the brain from pre-AD onset, revealed that AD develops as the brain changes over time [6]. Hence, treatment is speculated to be slow since symptoms have already emerged [7], and biomarker exploratory studies to start treatment early [8] and prevention studies have attracted attention.

Coffee is a popular beverage worldwide and has been consumed for many years because of its attractive flavor and physiological effects. It is also one of the best documented food items with epidemiological effects [9]. Epidemiological studies found that coffee consumption habits may reduce the risk of mild cognitive impairment and AD [10,11]. Coffee contains numerous phenolic compounds [coffee polyphenols (CPP)], such as chlorogenic acids (CGAs), and a single cup of coffee contains 70–350 mg of CGAs [12]. CGAs promote neuronal differentiation [13] and protect against Aβ-induced cell death by disaggregation of Aβ protein [14,15]. In addition, CGAs possess antioxidant activity, thereby improving temporary amnesia in mice [16]. These findings led us to hypothesize that CGAs could prevent memory and cognitive dysfunction induced by Aβ. However, whether CGAs have beneficial effects on AD is unknown.

APP/PS2 mice are double transgenic mice, overexpressing mutant forms of human Aβ precursor protein (hAPP) and human presenilin-2 (hPS2) [17,18]. Richards et al. reported that APP/PS2 mice exhibit AD-like impairments, such as amyloidosis, inflammation, impaired synaptic plasticity, and cognitive dysfunction [17].

In the present study, we used APP/PS2 mice as the model of AD to investigate whether memory and cognitive function could be maintained by treatment with CPP. These mice are a suitable model to investigate the therapy and prevention of AD because Aβ deposition was observed at 2–3 months of age in APP/PS2 mice. Thereafter, the cognitive decline was expressed at 4–5 months of age [18]. These behavioral and pathological changes in APP/PS2 mice are due to age-related cognitive dysfunction associated with amyloidosis.

## Materials and methods

### Preparation of CPP

Green coffee beans were extracted with hot water and spray dried. The extract was mixed with 52.4% ethanol solution, acid clay (MIZUKA ACE #600; Mizusawa Industrial Chemicals, Tokyo, Japan), and filter aid (SOLCA FLOC; JX Nippon Product Corporation, Tokyo, Japan) to obtain CPP obtaining slurry. The slurry was mixed with 52.4% ethanol solution and filtered with a diatomaceous earth filter. The filtrate was applied to activated charcoal column (SHIRASAGI WH2C; Osaka Gas Chemicals, Osaka, Japan) and cation-exchange resin column (SK1BH; Mitsubishi Chemical, Tokyo, Japan). The column processing solution was filtered using a 0.2-mm pore size membrane, and ethanol was removed with a rotary evaporator. The CPP-rich fraction was diluted with distilled water to yield a 3% (w/v) solution and centrifuged at 15 °C (1000×g, 60 min). Thereafter, the precipitate was lyophilized to analyze CPP composition through high-performance liquid chromatography. The total polyphenol content of the CPP was 48.3%. The composition of polyphenols was 7.0% 3-CQA, 7.5% 4-CQA, 17.8% 5-CQA, 1.3% 3-FQA, 1.5% 4-FQA, 3.9% 5-FQA, 3.4% 3,4-diCQA, 2.4% 3,5-diCQA, and 3.5% 4,5-diCQA. The CPP preparation contained no caffeine. The chemical structures of 5-CQA, 5-FQA, and 3,5-diCQA are shown in Fig 1.

**Fig 1.**
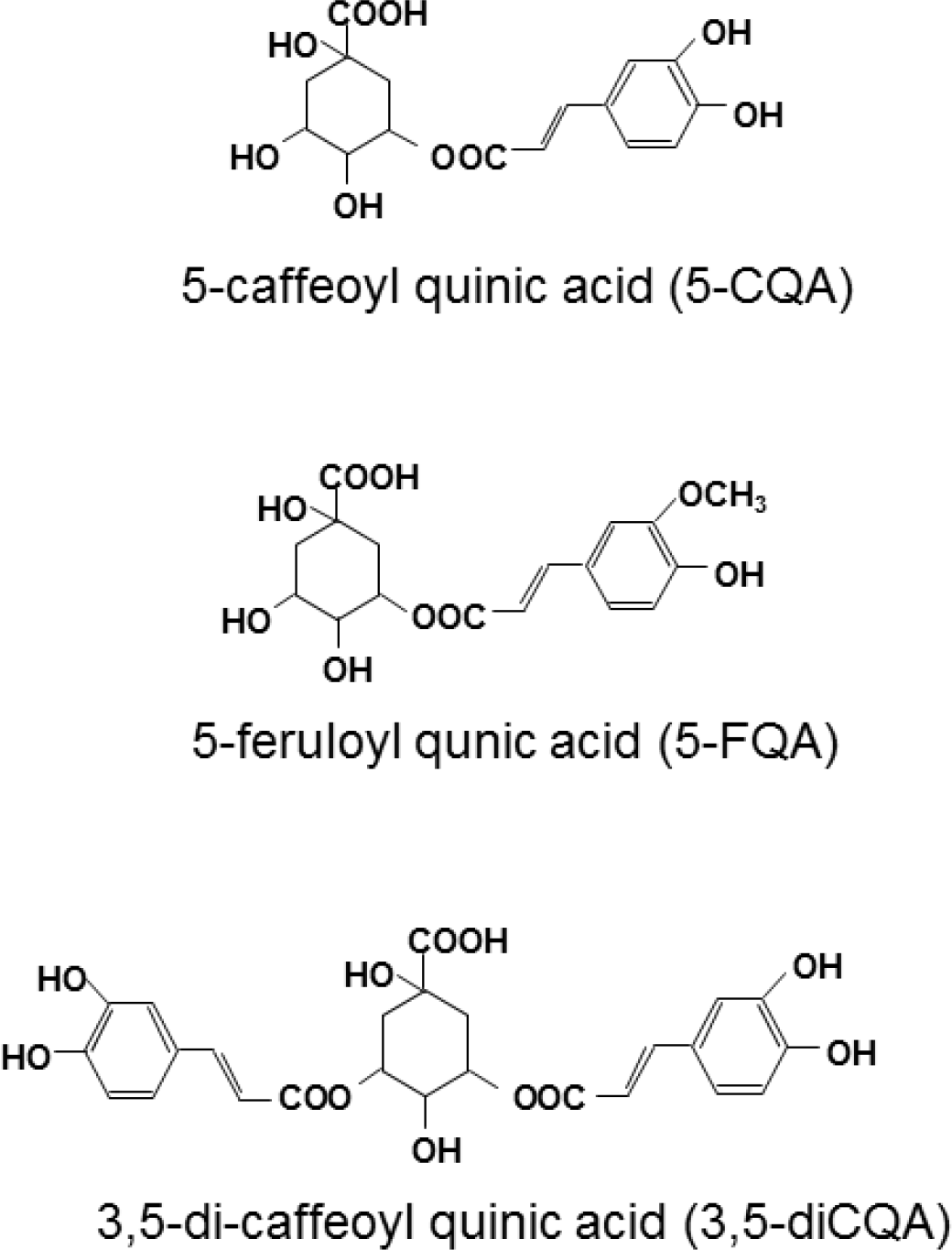
Structures of quinic acid derivatives.

### Animal and diets

The procedure of producing APP/PS2 double transgenic mice and wild-type (WT) littermates was determined using a method described by Toda et al. [18]. APPswe mice expressing hAPP gene mutant were purchased from Taconic Biosciences (Hudson, NY, USA). PS2M1 mice expressing hPS2 gene mutant were obtained from Oriental Yeast Co., Ltd. (Tokyo, Japan). APP/PS2 double transgenic mice were maintained by cross-breeding with APPswe male mice and PS2M1 female mice using in vitro fertilization and embryo transfer techniques. The mice genotype was confirmed through polymerase chain reaction analysis of tail-tip DNA.

Five-week-old male APP/PS2 mice and WT littermates were individually housed in a temperature- and humidity-controlled room (23 ± 3 °C, 55 ± 15% relative humidity) with a 12-h light/12-h dark cycle (lights on at 0600 h). The mice were divided into three groups (N = 12–15 mice/group). They were provided ad libitum access to water and either control or CPP diet. The control diet consisted of 10% (wt/wt) fat (corn oil), 20% casein, 61.5% potato starch, 4% cellulose, 3.5% vitamins, and 1% minerals. The CPP diet comprised the control diet supplemented with 1% CPP. Animals were maintained on these diets for 20 weeks. Individual body weights were recorded weekly, and food intake was measured every 3–4 days. Behavioral analysis was measured after 18 weeks as described below. After 20 weeks, the mice were anesthetized through isoflurane (Forane^®^ Abbott, Tokyo, Japan) inhalation. The brain was then transcardially perfused with 20 ml of saline, followed by 20 ml of 4% paraformaldehyde (PFA; Wako Pure Chemical, Osaka, Japan). The perfusate was maintained at 4 °C. After perfusion, the brain was dissected out, weighed, and stored in 4% PFA at 4 °C until analysis. All animal experiments were conducted in the Experimental Animal Facility of the Kao Tochigi Institute, and protocols were approved by the Kao Corporation Animal Care Committee (Protocol Number: F15045-0000). All surgery was performed under isoflurane anesthesia, and all efforts were made to minimize suffering.

### Behavioral analysis

Behavioral analysis was performed after 18 weeks in the following order: novel object recognition test, Morris water maze test, and step-through passive avoidance test.

#### Novel object recognition test

The novel object recognition test was performed in a plastic box (22 × 32 × 13 cm^3^). In the habituation trial on the first day, the mice were allowed to explore the empty test box for 10 min. On the following day, two objects (block) were placed in the test box. During the training trial, each mouse was placed in the test box and allowed to explore the objects for 10 min. Mice were returned to their home cage. After a 2-h training trial, one of the two objects (familiar) was replaced by a new one (novel), and test trial was carried out for 5 min. All sessions were video recorded. The exploration time spent by a mouse touching the object with its nose was measured. Data suggested that the exploration time spent by a mouse touching with the familiar or novel object is relative to the total exploration time during the test trial {[familiar or novel / (familiar + novel)] × 100}. The discrimination index was calculated as follows: {(novel – familiar) / (familiar + novel)}.

#### Morris water maze test

The Morris water maze pool, which had a diameter of 148 cm, contained water (17–18 °C) with a platform (10 cm diameter) that was submerged 2 cm beneath the water surface. Mice were first trained (4 days, 3 sessions/day) to find a hidden platform. The platform location remained invariable, and the entry point changed every trial. A trial was ended when mice stayed on the platform for 30 s or after 90 s. One day after day 4 of the training trial, the platform was removed and probe trial was carried out for 90 s. The entry point for the probe task was the reverse of the quadrant of the platform. The time taken for a mouse to swim to the previous quadrant of the platform was recorded. Performance was recorded and analyzed with the EthoVision XT (Version 7.0; Noldus Information Technology, Wageningen, Netherlands).

#### Step-through passive avoidance test

The step-through passive avoidance test chamber consisted of a light chamber (10 × 10 × 30 cm^3^) and a dark chamber (24 × 24.5 × 30 cm^3^), separated by a guillotine door (light and dark chamber; Nihon Bioresearch, Gifu, Japan). During the acquisition trial, each mouse was placed in the light chamber and allowed to search freely. After 10 s, the guillotine door was opened, allowing the mouse access to the dark chamber. The door closed after the mouse entered the dark chamber, and a 0.2-mA foot shock was administered to the mouse for 3 s (Shock scrambler; UNICOM, Tokyo, Japan). The latency time until the mouse entered the dark chamber was measured. Twenty-four hours after the acquisition trial, the mouse was placed in the light chamber for the test trial. The time taken for a mouse to enter the dark chamber was recorded.

#### Immunohistochemistry

To detect Aβ plaques, brain samples were fixed in 4% PFA, embedded in paraffin, and 4-µm-thick sections were obtained. Tissue sections were deparaffinized, treated with 90% formic acid for 5 min, and then incubated with 0.1% hydrogen peroxide in methanol to prevent endogenous peroxidation for 30 min. Subsequently, the tissue sections were incubated with the monoclonal antibody anti-human Aβ (#10323; Immuno-Biological Laboratories, Gunma, Japan, 1:200) at 25 °C for 1 h. The tissue sections were then incubated with HRP-conjugated streptavidin (Nichirei Biosciences, Tokyo, Japan) for 5 min, and diaminobenzidine substrate (DAKO, Tokyo, Japan) was used for color development. The images were acquired with a fluorescence microscope (BZ-X710; KEYENCE, Osaka, Japan), and quantitative image analysis was determined using BZ-II application (KEYENCE). Four slices for each mouse were used to quantify the mean average value of the selected regions. The Aβ deposition in the cerebral cortex and dorsal hippocampus was analyzed by the percentage of brain regions covered by Aβ immunoreactivity.

### RNA extraction and quantitative PCR (qPCR)

Total RNA was extracted using RNeasy Plus Universal Mini kit (Qiagen, Hiden, Germany). For real-time qPCR, cDNAs were synthesized with the High Capacity RNA-to-cDNA Kit (Applied Biosystems, Life Technologies, Forster City, CA, USA). qPCR assays were performed using an Applied Biosystems ViiA7 Real-time PCR system (Applied Biosystems). Commercially available polymerase chain reaction primers and FAM-labeled TaqMan probes (TaqMan Gene Expression assays; Applied Biosystems) were used for the assays. Expression of each gene was normalized against that of the gene encoding GAPDH, a housekeeping gene. The genes assessed in this study are listed in Supplemental S1 Table.

### Aβ fibrillization assays

Aβ fibrillization was measured with a commercially available thioflavin T (ThT) Aβ aggregation kit (Ana Spec, San Jose, CA, USA). The Aβ fibrillization assays were performed as described in the manufacturer’s procedure booklet. Aβ_1–42_ peptides were dissolved in dimethyl sulfoxide (DMSO) to prepare Aβ_1–42_ (2.5 mM) stock. 5-Caffeoylquinic acid (5-CQA) was dissolved in DMSO. Aβ_1–42_ solution (Ana Spec) was diluted to a final concentration of 50 µM. Aβ solution with ThT solution (200 µM; Ana Spec) and 5-CQA solution (100 µM; Cayman Chemical, Ann Arbor, MI, USA) was incubated at 37 °C in a black 96-well plate. The ThT fluorescence intensity was measured every 5 min for 90 min using the EnSight plate reader (Perkin Elmer, Waltham, MA, USA) at 440 nm (ex) and 484 (em).

### Aβ disaggregation assays

Aβ_1–42_ peptides (Peptide Institute, Osaka, Japan) were dissolved in DMSO to prepare Aβ_1–42_ (5 mM) stock. Subsequently, the Aβ_1–42_ stock was diluted with 50 mM Tris buffer to make Aβ_1–42_ (250 µM) solution. The Aβ_1–42_ solution was aggregated by incubation at 37 °C for 7–10 days. The aggregation and/or oligomerization state of Aβ_1–42_ solution was added with either DMSO or 5-CQA solution (1, 10, 100 µM, respectively) and incubated at 37 °C for 7 days. The final concentration of Aβ_1–42_ was 25 µM. The fibril formation of Aβ was measured using the ThT assay. Each incubated Aβ fibril formation was mixed with ThT solution (5 µM; Sigma Chemical Co, St. Louis, MO, USA). Fluorescence of Aβ bind to ThT was measured on the EnSight plate reader (Perkin Elmer) at 440 nm (ex) and 484 (em).

### Statistical analysis

Variables are expressed as the mean ± SEM. Statistical analysis was conducted using one-way analysis of variance (ANOVA), followed by Bonferroni’s post-hoc test or T-test (GraphPad Prism 6; GraphPad Software, La Jolla, CA, USA). Two-way repeated ANOVA, followed by Bonferroni’s post-hoc test, was used to assess changes over time and between groups (GraphPad Prism 6). Differences were considered significant when P < 0.05.

## Results

### Chronic treatment of CPP does not alter body and brain weight in APP/PS2 mice

During the CPP treatment, the general health conditions of APP/PS2 mice did not significantly change. The body weights of the WT, APP/PS2, and APP/PS2 + CPP groups were 46.18 ± 0.56, 48.03 ± 0.95, and 46.60 ± 0.00 g (N = 5 mice/group), respectively. The brain wet weights in the WT, APP/PS2, and APP/PS2 + CPP groups were 0.487 ± 0.003, 0.4989 ± 0.009, and 0.4986 ± 0.006 g (N = 5 mice/group), respectively. Both body and brain weights were not significantly different among the groups.

### Effect of CPP on the novel object recognition test

The effect of CPP on recognition memory was investigated using the novel object recognition test. In the training trial, each exploration time of the two objects was similar among three groups (data not shown).

The test trial was carried out after a 2-h training trial. In the WT and APP/PS2 + CPP groups, the percentage of novel object exploration time was increased (P < 0.001, Bonferroni’s post-hoc test) compared with the familiar object. However, the exploration time between familiar and novel objects did not significantly differ in the APP/PS2 group (Fig 2A).

When the exploration time of the objects was analyzed as a function of the discrimination index, the APP/PS2 group decreased (P < 0.001, Bonferroni’s post-hoc test) compared with the WT group (Fig 2B). By contrast, the discrimination index was increased in APP/PS2 mice treated with CPP (P < 0.001, Bonferroni’s post-hoc test), indicating that the recognition memory of CPP-treated APP/PS2 mice was improved (Fig 2B). The total exploration time during the test trial did not differ among the three groups (Fig 2C).

**Fig 2.**
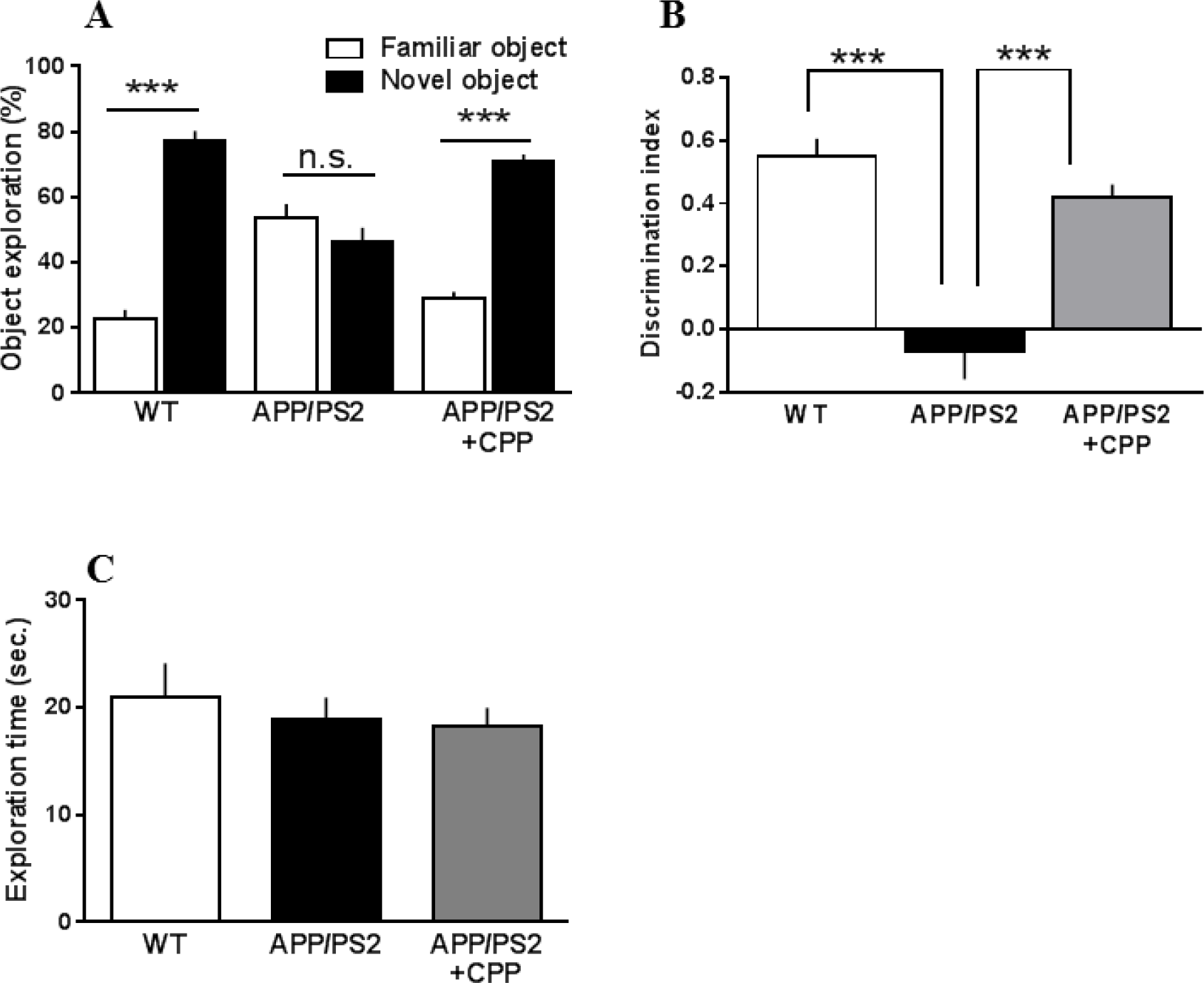
Effect of CPP on object recognition memory in the novel object recognition (NOR) test. (A) Percentage of exploration time for each object, (B) discrimination index, and (C) total object exploration time in the test session. The mice were fed experimental diets for 18 weeks prior to measurements. Values are the mean ± SEM of N = 12–15 mice/treatment. (A) ***: P < 0.001, familiar object vs. novel object (Bonferroni’s post-hoc test). (B), (C) ***: P < 0.001, vs. APP/PS2 (Bonferroni’s post-hoc test).

### Effect of CPP on the Morris water maze test

The effect of CPP on spatial learning and memory was investigated using the Morris water maze test. In the training trial, the escape latency time was significantly shorter (P < 0.05, Bonferroni’s post-hoc test) in the WT group than in the APP/PS2 group (Fig 3A). The escape latency time did not significantly differ between the WT and APP/PS2+CPP groups (Fig 3A). The swimming speed was the same among the three groups (Fig 3B).

Following the 4-day training trial, probe trials were demonstrated on the fifth day. The time swimming in the platform quadrant decreased (P < 0.01, Bonferroni’s post-hoc test) in the APP/PS2 group compared with the WT group (Fig 3C), whereas improved in the APP/PS2+CPP group compared with the APP/PS2 group (Fig 3C). This improvement was comparable with the WT group.

**Fig 3.**
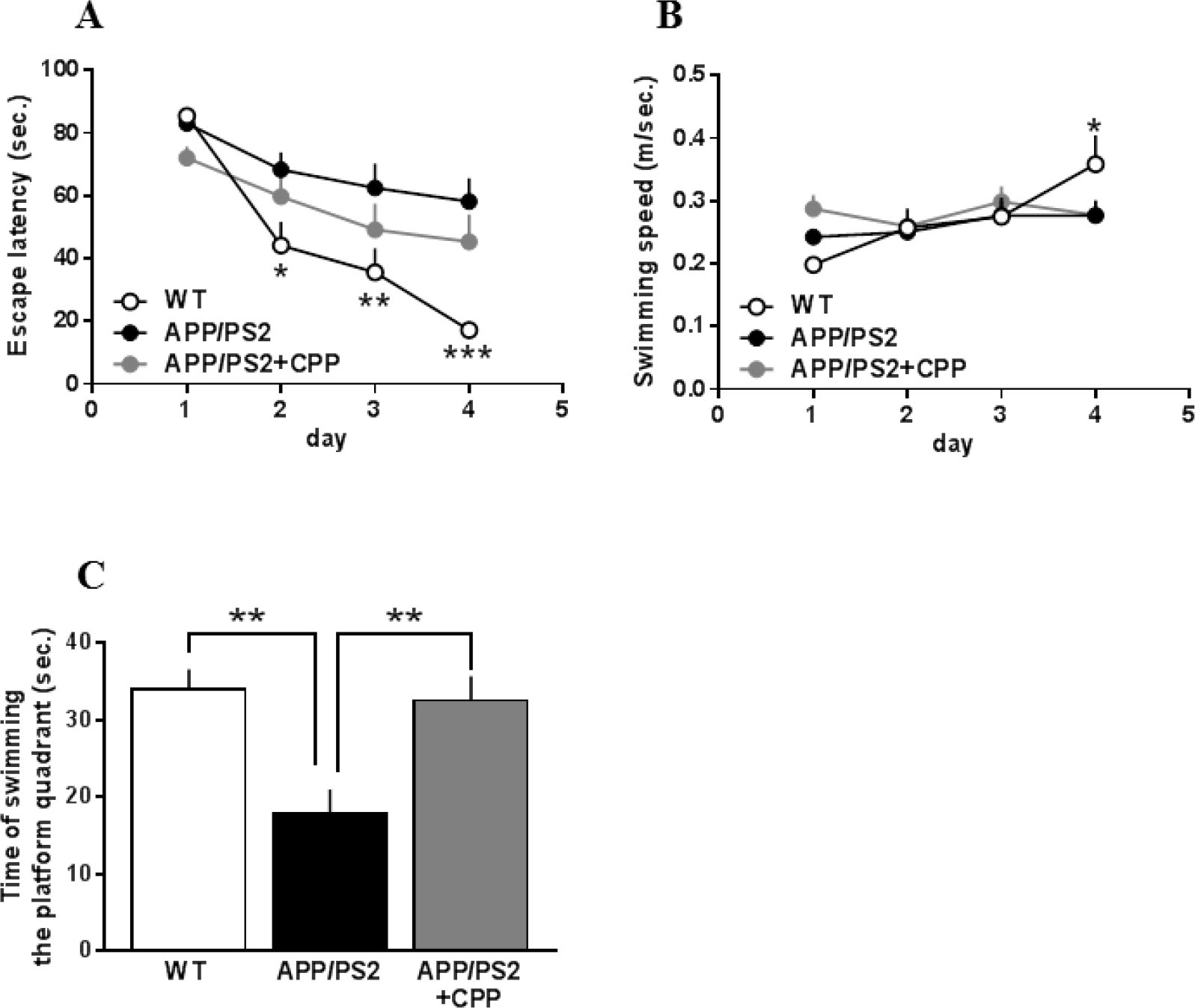
Effect of CPP on spatial learning and memory in the Morris water maze test. (A) Escape latency and (B) swimming speed during training task. (C) Time required swimming the target quadrant in the probe test. The mice were fed experimental diets for 19 weeks prior to measurements. Values are the mean ± SEM of N = 12–15 mice/treatment. *: P < 0.05, **: P < 0.01, ***: P < 0.001, vs. APP/PS2 (Bonferroni’s post-hoc test).

### Effect of CPP on the step-through passive avoidance test

The effect of CPP on long-term memory was investigated using the step-through passive avoidance test. In the acquisition trial, the latency time was similar in all groups (Fig 3). In the test trial after the 24-hour acquisition trial, the latency time was significantly longer (P < 0.05, Bonferroni’s post-hoc test) in the WT group than in the APP/PS2 group (Fig 4). The latency time tended (P = 0.10, Bonferroni’s post-hoc test) to be improved in the APP/PS2 + CPP group than in the APP/PS2 group (Fig 4).

**Fig 4.**
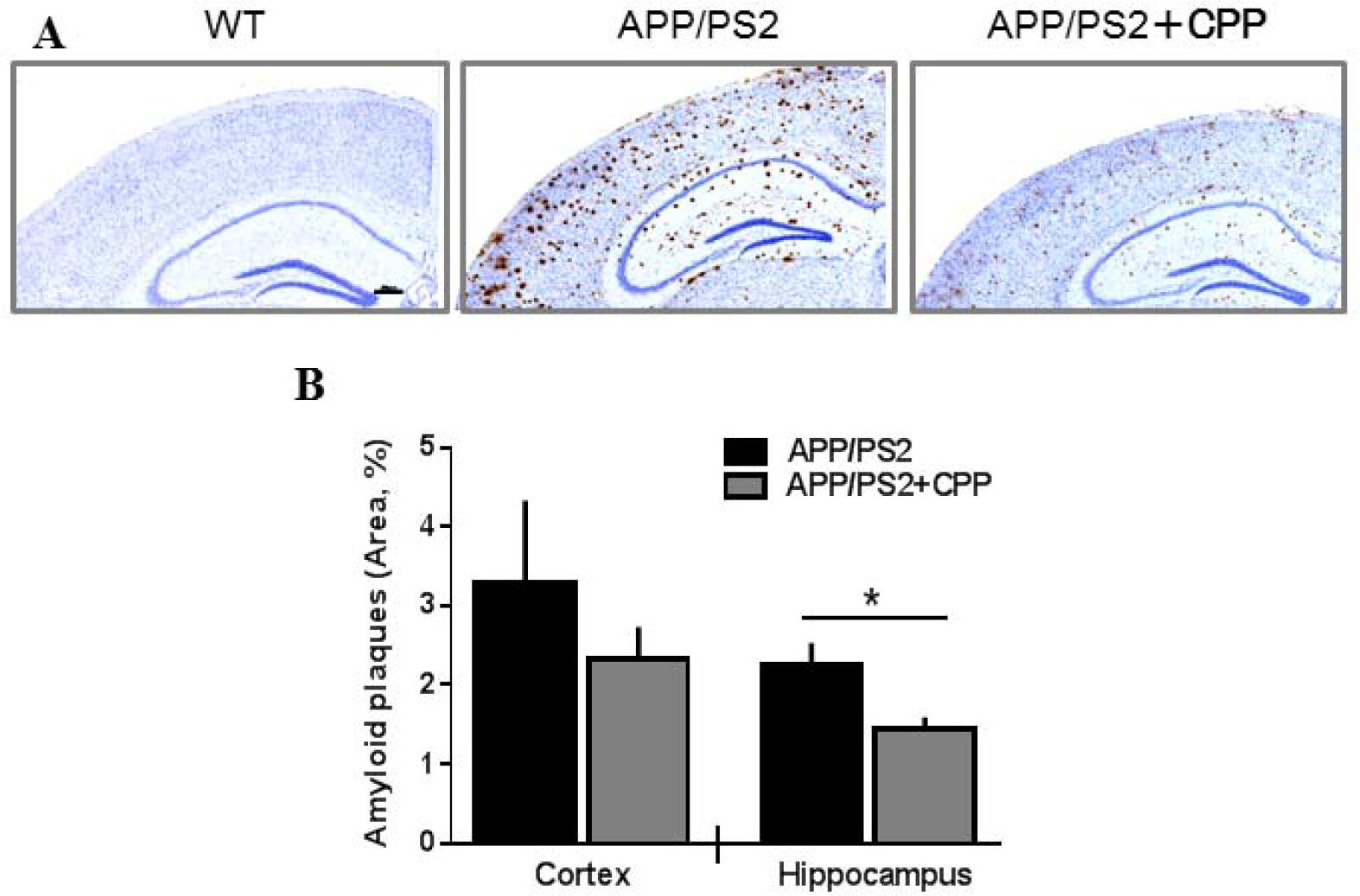
Effect of CPP on the step-through passive avoidance test. Mice were fed experimental diets for 21 weeks prior to measurements. Values are the mean ± SEM of N = 12–15 mice/treatment. *: P < 0.05, vs. APP/PS2 (Bonferroni’s post-hoc test).

### Effect of CPP on Aβ deposition in the brain

Aβ plaque increased in the cortex and hippocampus of the APP/PS2 group compared with the WT group (Fig 5A), whereas it decreased in those of the APP/PS2 + CPP group compared with the APP/PS2 group (Fig 5A). The area of Aβ plaque in the hippocampus was significantly decreased (P < 0.05, T-test) in the APP/PS2 + CPP group compared with that in the APP/PS2 group (Fig 5B).

**Fig 5. Effect of CPP on Aβ pathology in the brains of APP/PS2 mice**.(A) Immunohistochemistry of Aβ plaques in the brain slices of WT, APP/PS2, and APP/PS2+CPP mice. The black bar is equal to 200 µm for 40× magnification. (B) Quantification of Aβ plaques is expressed with bar graphs of mean percentage of area. Values are the mean ± SEM of N = 6–8 mice/treatment. *: P < 0.05, vs. APP/PS2 (student t-test).

### Effect of CPP on gene expression in the mouse brain

APP/PS2 mice showed significantly higher mRNA expression of CatB, NOX2, and p22phox than wild type mice in both the hippocampus and cortex (Table 1). Expression of proinflammatory genes, such as CD68, F4/80, IL-1β, and IL-6, was also significantly increased in APP/PS2 mice than in control mice in either the hippocampus and cortex. APP/PS2 mice exhibited significantly higher gene expression of glial cell markers, such as A1, Iba1, and GFAP, than the control mice in either the cortex or hippocampus. CPP-fed APP/PS2 mice had slightly lower mRNA expression of mouse APP in the cortex, but not in the hippocampus, than CPP-non-fed APP/PS2 mice (Table 1). The expression of other genes tested did not differ between CPP-fed APP/PS2 and control mice (Table 1).

**Table 1.** Effects of CPP on mRNA expression in the mouse brain

|  |  |  | WT | APP/PS2 | APP/PS2+CPP |
| --- | --- | --- | --- | --- | --- |
| **Amyloid metabolism** |  |  |  |  |  |
| ADAM10 | Hippocampus |  | 1.01 ± 0.03 | 1.00 ± 0.01 | 0.97 ± 0.06 |
|  | Cortex |  | 1.07 ± 0.02* | 1.00 ± 0.02 | 1.00 ± 0.02 |
| APP (human) | Hippocampus |  | N.T. | 1.00 ± 0.02 | 1.01 ± 0.06 |
|  | Cortex |  | N.T. | 1.00 ± 0.02 | 0.96 ± 0.03 |
| APP (mouse) | Hippocampus |  | 1.01 ± 0.02 | 1.00 ± 0.02 | 0.87 ± 0.05** |
|  | Cortex |  | 1.10 ± 0.01*** | 1.00 ± 0.02 | 0.95 ± 0.02 |
| BACE1 | Hippocampus |  | 0.96 ± 0.02 | 1.00 ± 0.02 | 0.94 ± 0.05 |
|  | Cortex |  | 1.16 ± 0.02** | 1.00 ± 0.02 | 1.10 ± 0.05 |
| CatB | Hippocampus |  | 0.74 ± 0.01*** | 1.00 ± 0.03 | 0.97 ± 0.06 |
|  | Cortex |  | 0.77 ± 0.04*** | 1.00 ± 0.02 | 1.02 ± 0.03 |
| **Inflammation** |  |  |  |  |  |
| CD68 | Hippocampus |  | 0.71 ± 0.03** | 1.00 ± 0.04 | 0.96 ± 0.09 |
|  | Cortex |  | 0.60 ± 0.03** | 1.00 ± 0.07 | 0.89 ± 0.13 |
| F4/80 | Hippocampus |  | 0.49 ± 0.05*** | 1.00 ± 0.03 | 0.97 ± 0.11 |
|  | Cortex |  | 0.46 ± 0.03*** | 1.00 ± 0.05 | 0.84 ± 0.10 |
| IL-1β | Hippocampus |  | 0.26 ± 0.03** | 1.00 ± 0.10 | 1.07 ± 0.20 |
|  | Cortex |  | 0.37 ± 0.06** | 1.00 ± 0.08 | 1.10 ± 0.19 |
| IL-6 | Hippocampus |  | 0.42 ± 0.06*** | 1.00 ± 0.11 | 1.10 ± 0.07 |
|  | Cortex |  | 0.44 ± 0.06** | 1.00 ± 0.11 | 1.28 ± 0.13 |
| iNOS | Hippocampus |  | 1.01 ± 0.09 | 1.00 ± 0.03 | 1.09 ± 0.13 |
|  | Cortex |  | 0.93 ± 0.05 | 1.00 ± 0.05 | 1.04 ± 0.17 |
| **Redox regulation** |  |  |  |  |  |
| NOX2 | Hippocampus |  | 0.40 ± 0.09*** | 1.00 ± 0.06 | 1.15 ± 0.11 |
|  | Cortex |  | 0.27 ± 0.03*** | 1.00 ± 0.10 | 1.04 ± 0.14 |
| p22phox | Hippocampus |  | 0.49 ± 0.04*** | 1.00 ± 0.04 | 1.10 ± 0.12 |
|  | Cortex |  | 0.32 ± 0.01*** | 1.00 ± 0.03 | 1.09 ± 0.07 |
| **Neuroendocrine cells** |  |  |  |  |  |
| Synaptophysin | Hippocampus |  | 0.99 ± 0.02 | 1.00 ± 0.01 | 0.93 ± 0.05 |
|  | Cortex |  | 1.09 ± 0.02 | 1.00 ± 0.02 | 1.06 ± 0.06 |
| **Glial cells** |  |  |  |  |  |
| A1 | Hippocampus |  | 0.17 ± 0.01*** | 1.00 ± 0.07 | 1.08 ± 0.10 |
|  | Cortex |  | 0.13 ± 0.01*** | 1.00 ± 0.06 | 1.12 ± 0.05 |
| GFAP | Hippocampus |  | 0.30 ± 0.02*** | 1.00 ± 0.06 | 1.08 ± 0.13 |
|  | Cortex |  | 0.12 ± 0.01*** | 1.00 ± 0.04 | 1.15 ± 0.09 |
| Iba-1 | Hippocampus |  | 0.56 ± 0.01*** | 1.00 ± 0.05 | 1.01 ± 0.10 |
|  | Cortex |  | 0.56 ± 0.03*** | 1.00 ± 0.06 | 1.05 ± 0.06 |

Values are relative gene expression (means ± SEM) of 6-8 mice (APP/PS = 1). Statistical analysis was conducted using one-way analysis of variance, followed by Bonferroni's post-hoc test or T-test (GraphPad Prism 6; GraphPad Software, La Jolla, CA, USA). Differences were considered significant when P < 0.05.
\*: $P < 0.05$ , \*\*: $P < 0.01$ vs. APP/PS2 group. N.T., not tested; A1, B cell leukemia/lymphoma 2 related protein A1a; ADAM10, a disintegrin and metallopeptidase domain 10; BACE1, beta-site APP cleaving enzyme 1; CatB, cathepsin B; F4/80, EGF-like module-containing mucin-like hormone receptor-like 1; GFAP, glial fibrillary acidic protein; Iba-1, allograft inflammatory factor 1; iNOS, inducible nitric oxide synthase.

### Effect of 5-CQA on Aβ formation

The fluorescence intensity of Aβ alone increased immediately and reached a plateau after a 60-min incubation. The presence of 5-CQA, the main component of CPP, did not affect the fluorescence intensity, suggesting that 5-CQA did not inhibit Aβ fibrillization (Fig 6A).

The aggregation and/or oligomerization state of Aβ was disaggregated with increasing 5-CQA dosage, and it was significantly lower (P < 0.05, Bonferroni’s post-hoc test) in the 5-CQA (10 and 100 µM) than in the without 5-CQA (Fig 6B).

**Fig 6.**
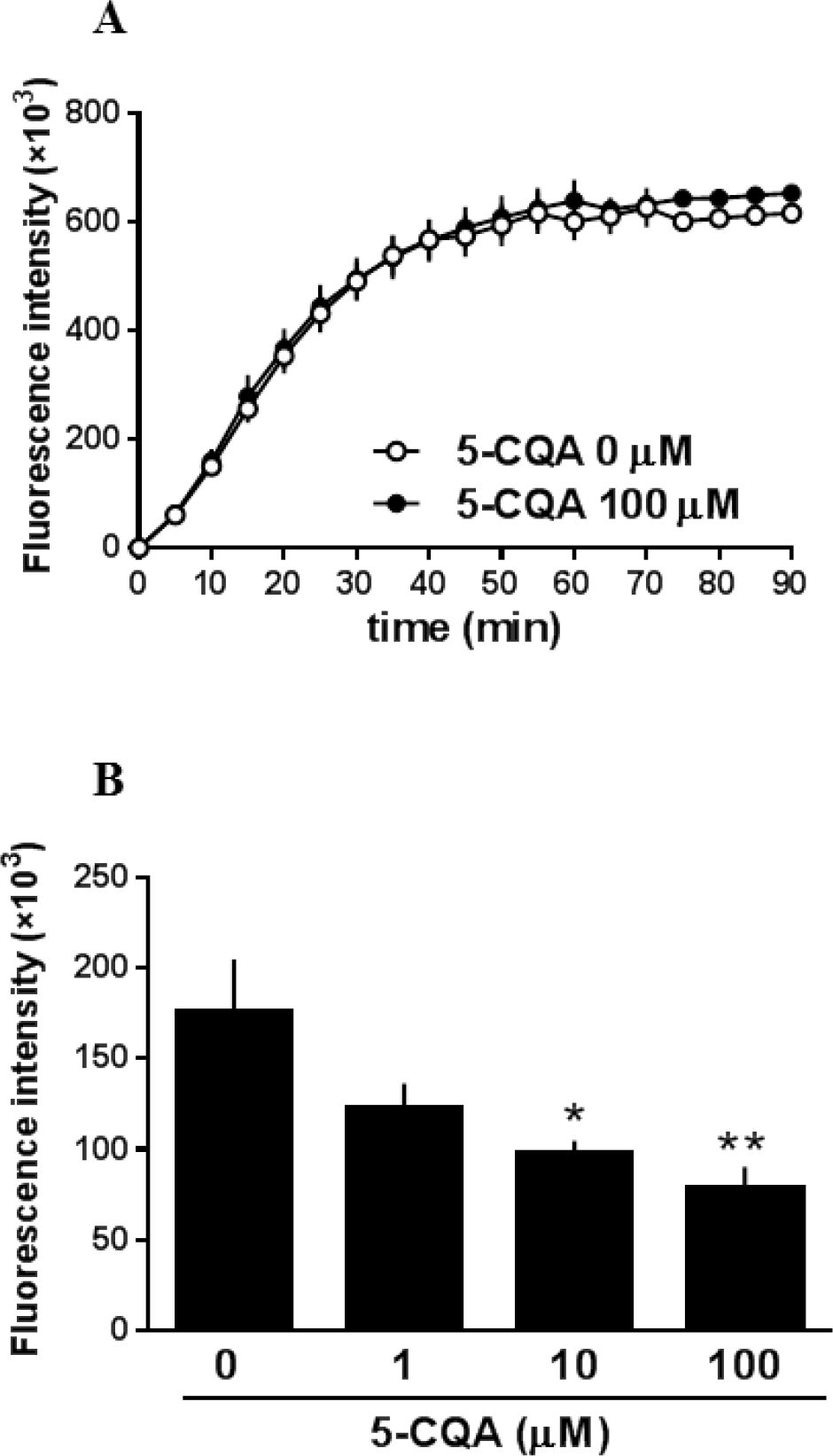
Effect of 5-CQA on Aβ fibril formation in vitro. The formation of Aβ_1–42_ was measured using the ThT fluorescence assays. (A) Aβ_1–42_ solution (200 µM) was incubated with 5-CQA (100 µM). The time course of changes in the fluorescence intensity was measured. Fluorescence intensity is expressed as Aβ fibrillization. (B) The aggregation and/or oligomerization state of Aβ_1–42_ (25 µM) was incubated with either DMSO or 5-CQA (1, 10, 100 µM) for 7 days before the assay. Fluorescence intensity is expressed as Aβ disaggregation. Values are the mean ± SEM of N = 5–8. *: P < 0.05, **: P < 0.01, vs. without 5-CQA (Bonferroni’s post-hoc test).

## Discussion

In general, the effectiveness of coffee for health is owing to the abundant polyphenols in coffee beans typified by CGAs. In this study, we found that chronic consumption of CPP improved memory and cognitive function and reduced Aβ pathology in APP/PS2 mice. We also demonstrated that the CPP may modulate AD phenotypes by promoting disaggregation of fibril Aβ species into Aβ peptide in the brain.

The CPP-treated mice improved memory and cognitive function because of three behavioral analyses. The novel object recognition test is based on the spontaneous tendency of rodents to explore a novel object without requiring reward or penalty. This test measures visual recognition memory, the spatial and temporal context of object recognition, which is supported by interactions between the cortex and the hippocampus [19]. The Morris water maze test investigates spatial learning and memory. This test involves the hippocampus, cerebral cortex, and striatum [20]. The step-through passive avoidance test evaluates long-term memory of learned avoidance behavior by electric foot shock. These behavioral analyses may influence not only cognitive function, but also motivation or motor function [21]. Improvement in the performance of learning and memory tasks in CPP-treated mice is unlikely due to the changes in activity, motivation, or motor function. The total object exploration time in the novel object recognition test, the swimming speed in the Morris water maze test, and the latency time during acquisition trial in the step-through passive avoidance test in CPP-treated mice did not differ from those in CPP-non-treated mice. These results suggest that exploratory activity and motor function, that is swimming ability, did not change. Therefore, the improved behavioral performance in CPP-treated mice is probably owing to enhanced learning and memory. Several studies suggested that hippocampal damage causes impaired performance in behavioral tests [20, 22, 23]. The CPP-treated mice showed significantly decreased Aβ plaque in the hippocampus than the CPP-non-treated mice. These findings suggest that the CPP protected against Aβ toxicity, particularly in the hippocampus, and improved the three behavioral test performances.

In this study, dietary CPP significantly decreased Aβ deposit in the brain of APP/PS2 mice without changing Aβ metabolism-related gene expression. To study the underlying mechanism for the reduction of Aβ deposit by dietary CPP, we investigated the effects of CPP components on the aggregation and disaggregation of Aβ-protein.

CGAs possess inhibitory effect on Aβ-protein aggregation [14]. Therefore, the influence of 5-CQA, a major component of CPP, on Aβ fibrils was investigated using the ThT assay, and Aβ fibrils were disaggregated, which did not affect Aβ fibril formation. Ono et al. reported that grape seed-derived polyphenols with phenolic groups like CGA disaggregate Aβ fibrils and reduce the cytotoxicity of Aβ fibrils [24]. Wei et al. reported that CGA has a neuroprotective effect on the cytotoxicity of Aβ [15]. These studies suggest that CGA may disaggregate Aβ fibrils and could possibly reduce cytotoxicity. Conventionally, Aβ fibrils that accumulate as cerebral amyloid are thought to exert neurotoxicity. However, in recent years, oligomers, the intermediate stages of amyloid β protein aggregation, have been reported to be the most toxic, and oligomer reduction may be effective for treating AD [25]. Therefore, investigating the influence of CGA on the structure of amyloid β protein and the formation and degradation of oligomers in the future is necessary.

Aβ deposition is observed in APP/PS2 mice from 2 to 3 months of age and it increases with age; cognitive function decreases after 4–5 months of age [17, 18]. Fontata and colleagues reported that synaptic excitability of the hippocampus changes from 3 months of age in APP/PS2 mice [26]. In addition, the increase in Aβ deposition in humans begins 15–20 years before AD onset [6]. Protecting the brain from disorders caused by Aβ and preventing the onset of AD through early intervention are important. The cognitive dysfunction is not improved when it develops in AD only by removal of Aβ in the brain [27]. In this study, CGA was ingested by 5-week-old mice before the changes in Aβ deposition, cognitive function, and synaptic function occurred, and the preventive effect of CGA on AD onset was clarified. Therefore, in this study, we could demonstrate the effect of CGA. Further studies on the analysis of the detailed mechanism of CGA on brain functions, and its effectiveness on Aβ deposition, which has already occurred as well as cognitive decline, are necessary. CGA does not only have an effect in reducing Aβ deposition, but also has inhibitory actions on antioxidant and AChE activities [16]. Thus, these findings suggest that CGA has various effects to improve cognitive dysfunction and also therapeutic effect in AD.

## Conclusion

We demonstrated that CPP treatment significantly attenuated recognition memory, spatial learning and memory, and long-term memory activity decline, as well as alleviated Aβ plaque deposition. Consequently, CPP provided a protective effect against AD progression through Aβ. These findings may shed light on CPP as an effective therapeutic and prophylactic agent for cognitive deficits by AD.

## Acknowledgments

We thank Tatsuya Kusaura and Yukiteru Sugiyama for preparing the CPP. We thank Yukie Kawatani for technical assistance.

## Author Contributions

**Conceptualization:**KI KM AS NO.

**Formal analysis:**KI MY.

**Investigation:**KI MY.

**Methodology:**KI MY KM.

**Project administration:**NO.

**Supervision:**KM AS NO.

**Visualization:**KI MY.

**Writing-original draft:**KI.

**Writing-review & editing:**KM AS.

## Supporting information

**S1 Table. Taqman probes**.

